# Vcfexpress: flexible, rapid user-expressions to filter and format VCFs

**DOI:** 10.1101/2024.11.05.622129

**Authors:** Brent S. Pedersen, Aaron R. Quinlan

## Abstract

**Motivation:** Variant Call Format (VCF) files are the standard output format for various software tools that identify genetic variation from DNA sequencing experiments. Downstream analyses require the ability to query, filter, and modify them simply and efficiently. Several tools are available to perform these operations from the command line, including BCFTools, vembrane, slivar, and others.

**Results:** Here, we introduce vcfexpress, a new, high-performance toolset for the analysis of VCF files, written in the Rust programming language. It is nearly as fast as BCFTools, but adds functionality to execute user expressions in the lua programming language for precise filtering and reporting of variants from a VCF or BCF file. We demonstrate performance and flexibility by comparing vcfexpress to other tools using the vembrane benchmark.

**Availability:** vcfexpress is available under the MIT license at https://github.com/brentp/vcfexpress

**Supplementary information:** Supplementary data are available at *Bioinformatics* online.

## 1 Introduction

Files in Variant Call Format(Danecek *et al*., 2011) (VCF) are the starting point of many genetic analyses. The VCF format can represent diverse metadata detailing each genetic variant, as well as the distinct genotypes and haplotypes observed for each variant. While it provides a comprehensive container for all relevant data, the complexity of the format poses challenges for extracting the information germane to diverse downstream analyses. Therefore, it is critical to have the capability to easily and rapidly filter, query, and format these files. There are several tools available to perform these operations; notably BCFTools(Danecek *et al*., 2021), slivar(Pedersen *et al*., 2021), vembrane(Hartmann *et al*., 2023), SnpSift(Cingolani, Patel, *et al*., 2012), and bio-vcf(Garrison *et al*., 2022). Here we introduce vcfexpress, which we provides a powerful combination of speed, safety, functionality, expressiveness, and ease of use. In contrast, slivar is best for expressions that filter variants based on sample and family attributes and patterns. Furthermore, other tools like vembrane and BCFTools are similar in intent to vcfexpress; however, vcfexpress expands on their functionality by allowing complete control over the filtering expressions and data selection from a VCF, while retaining high processing performance.

## 2 Methods

### 2.1 Implementation

Vcfexpress is implemented in the rust programming language using rust-htslib (https://docs.rs/rust-htslib/latest/rust_htslib/) which wraps the HTSlib(Bonfield *et al*., 2021) C library. Internally, vcfexpress relies on a lua wrapper of rust-htslib, allowing users to pass custom lua expressions (specifically, luau) to vcfexpress at runtime. This functionality represents an important advance in vcfexpress, as these expressions provide unique analytical power and flexibility. In addition to filtering VCF files in search of variants that meet diverse, user-defined criteria, these filters can be used to modify and add VCF fields. Therefore, users can employ vcfexpress to add new annotations to empower downstream analyses. Because of the software’s processing speed, we were able to implement support for parsing SnpEff(Cingolani, Platts, *et al*., 2012) and VEP(McLaren *et al*., 2016) functional annotations with a user-defined lua script rather than as specialized built-in code; this functionality demonstrates the flexibility of our approach. Because we use the luau variant of lua, expressions can be run in a sandboxed environment (https://luau.org/sandbox) allowing safe execution of untrusted user code and expressions. Another unique use-case is a template argument that can specify the output format (e.g., BED, BEDGRAPH) of the filtered and/or modified VCF.

### 2.2 Usage

#### Pedersen et al

Vcfexpress is called from the command line with a user-provided filter expression that is applied to each variant that returns a boolean indicating whether the variant should be written to the output based on the results of the filter expression. In addition, the user can provide a template string that indicates how to format the output of the filtered variants. If this is not given, then the output is formatted as VCF (or BCF depending on the suffix of the output file). The following example illustrates using the vcfexpress “filter” function to exclusively report records if all samples have a sequencing depth greater than 10 and the AN info field is above 100. The template argument specifies that vcfexpress should output a BED file of all passing variants:

~~~
vcfexpress filter \
 -e ‘return all(function (dp) return dp > 10
end, variant:format(“DP”)) \
             and variant:info(‘AN’) > 100’ \
   --template ‘{variant.chrom}\t{variant.start}\
t{variant.stop}’ \
  -o all-high-dp.bed $input_vcf
~~~

To our knowledge, such template functionality is not available in other tools, and it provides important analytical flexibility, especially for VCF files with many samples. The next example shows how the user can modify the BCF input to have a new INFO field called *HIGH_IMPACT*. This new field is populated using information extracted from the CSQ field and written to a new BCF as output.

~~~
vcfexpress filter \
  -s ‘HIGH_IMPACT=return
CSQS.new(variant:info(‘vep’), desc):any(
          function(c) return c.IMPACT ==
‘HIGH’ end)’ \
      -p scripts/csq.lua \
      -p scripts/high_impact_header.lua \
      -o high_impact.bcf input.bcf
~~~

This example uses *scripts/high_impact_header*.*lua*, which contains the code to update the VCF header:

~~~
header:add_info({ID=“HIGH_IMPACT”, Number=0,
Description=““, Type=“Flag”})
desc =
parse_description(header:info_get(“vep”).Descrip
tion)
~~~

We emphasize that these are mere examples of the vast analytical flexibility provided by the lua filtering language and full usage is documented on the github page for the tool.

## 3 Results

We assessed vcfexpress’s performance using the vembrane benchmark to compare the speed of several tools on a set of filtering tasks ranging from simple INFO field parsing to per-sample attributes to parsing the consequence (CSQ) fields. Each tool has specific strengths and utilities; these are documented nicely in the vembrane paper(Hartmann *et al*., 2023). Generally, BCFTools has been the fastest; however, we find that vcfexpress has similar speed, while supporting advanced filtering logic and VCF and BCF formats (**Figure 1**).

**Figure 1.**
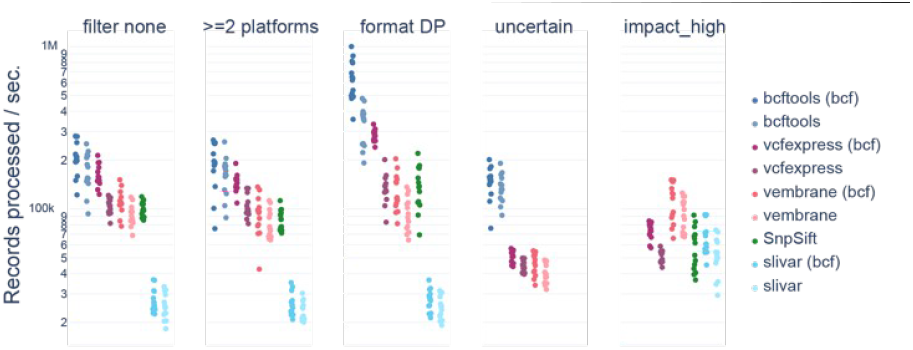
Speed of tools across a range of filters. Here we compare five tools in filtering VCF and (where possible) BCF. The filters are: *filter_none* (remove all records); *>=2 platforms* (filter on numeric INFO field); *format DP* (require at least 1 sample to have high depth); *uncertain* (check the VEP consequence field for “uncertain_significance”); *impact_high* (check if any CSQ has high impact). Note that vcfexpress is competitive with bcftools for speed while offering extreme flexibility.

## Summary

We have introduced vcfexpress which supports user expressions for filtering, formatting, and modifying records in VCF and BCF files. We show that it is on par with the fastest tool and expands upon existing variant filtering functionality. The lua expressions are extremely flexible–offering the full power of the lua language while maintaining high processing performance. We therefore expect that vcfexpress will be a useful tool in this space.

## Acknowledgments

We acknowledge Tom Sasani for their helpful discussions and assistance with editing the manuscript.

